# Seasonal life-history trade-offs differ between plants derived from seed versus rhizomes in *Saponaria officinalis*

**DOI:** 10.1101/2025.10.28.685121

**Authors:** Alyson C. Van Natto, Jannice Friedman

## Abstract

Many perennial plants reproduce both sexually through seeds and asexually through vegetative clones, and these contrasting origins can shape subsequent life-history strategies. Because clonal propagules begin with greater parental resource provisioning, whereas seedlings establish independently, these alternative origins create natural variation in early resource availability and developmental trajectories. This variation provides an opportunity to test predictions from life-history theory: while investment in current reproduction is expected to trade off with future growth or survival, such costs are often difficult to detect unless reproductive effort varies substantially among individuals or is experimentally manipulated. In this study, we compared growth, reproduction, and survival of seed- and rhizome-origin individuals of the invasive perennial *Saponaria officinalis* across three years in a common garden. We also manipulated reproductive effort in a subset of plants through pollen supplementation and tracked seasonal changes in flowering and seed production. Seedlings initially grew faster and made more branches and flowers than rhizome-origin plants, but by the second year rhizomes were larger and had more flowers. Seedlings also exhibited far worse survival than rhizomes. Across the flowering season, seed number per capsule declined, consistent with both increasing competition among flowers and declining resource availability. Pollen-supplemented seedlings had the most seeds per capsule early in the season and were the only group to suffer overwinter mortality, indicating a survival cost of elevated early reproductive investment. Overall, our findings show that reproductive origin influences allocation to growth, reproduction, and survival, with consequences for the establishment and spread of this invasive weed.

## Introduction

Most perennial plants can reproduce both sexually and asexually (Klimeš et al., 1997; Barrett, 2015), with the success of each mode shaped by selection that ultimately determine a plant’s life-history strategy. Sexual and asexual reproduction differ in their costs and benefits, and the likelihood of successful establishment for each type of propagule depends on ecological conditions. Seeds, arising from sexual reproduction, are more likely to colonize new habitats than clonal fragments because seeds often disperse more readily. However, in established populations, recruitment from seeds can be infrequent (Eriksson, 1992). Because seedlings rely on stored seed reserves for germination and early growth, they are more vulnerable to environmental unpredictability and competition (Fenner, 1987; Lei, 2010). Unless the habitat is entirely open, seedlings face competition, whether it’s from the parent plant, germinating siblings or other seedlings, or existing vegetation (Fenner, 1987). In contrast to seeds, clonal propagules do not face the same establishment risks, as they arise from an existing individual and can immediately benefit from resource sharing, reducing early mortality (Lei, 2010). Clonal propagules are often larger, more robust, and competitive than seeds because they are better provisioned with maternal resources. Competition among clones may occur but is more likely at high densities or under resource limitation (Rautainen et al., 2004). Because the path to establishment is inherently different between seedlings and clonal propagules, we may expect them to diverge in the timing and allocation to key life cycle events.

Among clonal propagules, ramets—genetically identical offspring that arise through vegetative offshoots—represent one of the most common forms of asexual reproduction in perennial plants. Several studies have explored how the growth and survival of ramets compare to those of sexually derived seedlings, but results have been mixed. For example, in a greenhouse common garden experiment examining growth metrics of *Spartium junceum*, seedlings had greater dry mass than ramets (Nilsen and Semones, 1997). In contrast, after one growing season, ramets of four grassland species (*Poa trivialis, Poa annua, Agrostis stolonifera*, and *Festuca rubra*) were 18 times the dry weight of seedlings and had higher survival rates (Howe & Snaydon, 1986). Similarly, ramets of *Imperata cylindrica* were taller and had greater survival than seedlings at low densities, however ramet and seedling performance was similar at high densities (Kushwaha et al., 1983). While these studies demonstrate differences between ramets and seedlings, all were limited to a single growing season, and offer little insight into multi-year strategies that are relevant for perennial plants. This short-term focus also means they did not examine reproductive allocation or its potential trade-offs with future growth and survival.

Life-history theory predicts that current reproduction should trade off with growth due to limited resources (Stearns 1989). There could be subsequent trade-offs between current reproduction and future survival, representing costs of reproduction (Williams 1966; Dorken 2025). For example, in *Helianthemum squamatum* (Aragón et al., 2009) and *Carex secalina* (Bogdanowicz et al., 2011), reproduction in one year negatively affected future survival and fecundity. In their recent comprehensive review, Dorken (2025) find that overall 83% of studies identify costs of reproduction. However, studies that failed to identify costs were disproportionately conducted in natural settings (89%) where trade-offs are notoriously difficult to detect. The challenge in this case is that plant resource status, size, and age are not controlled at the start of the experiment, so that expected trade-offs between life history components are masked. The result is that the difference in initial photosynthetic capacity, resource acquisition, or resource availability, leads to positive correlations between traits that are expected to vary negatively (Obeso, 2002; van Noordwijk & de Jong 1986). Other reasons for the general absence of costs are physiological: plants may reabsorb resources from flowers or aborted fruits and reallocate them (Ashman, 1994); or photosynthesis in floral structures may offset their cost (Reekie & Bazzaz, 1987). These compensatory mechanisms may prevent trade-offs from being detected so that experimentally imposing environmental stress can expose otherwise hidden reproductive costs (Sandvik, 2001; Hamann et al., 2021). Another approach to detecting trade-offs is to experimentally manipulate reproductive effort, either by removing reproductive structures or by supplementing pollen, to determine how changes in reproductive effort affects survival, and future growth and reproduction.

Many studies have examined whether seed set is primarily limited by pollen availability or resource constraints. Typically, this is tested with pollen supplementation experiments: if supplemental pollen increases seed set then pollen limitation is the primary constraint; whereas if seed set does not increase then resources are likely the limiting factor. Across species, the strength of pollen limitation varies, with some studies finding strong pollen limitation (e.g., Larson & Barrett, 1999; Knight, 2003), while others report mixed evidence or no pollen limitation (e.g., Totland, 1997; Eckert & Schaefer, 1998). Although studies using supplemental pollen have contributed significantly to our understanding of reproductive constraints, they also have limitations. For example, most studies only supplement a fraction of flowers, which can lead to the reallocation of resources to pollinated flowers, potentially concealing resource costs for full seed set (Ashman et al., 2004). To better address whether plants have the resources to fully provision all the ovules in their flowers, most or all flowers should be supplemented with pollen (Ashman et al., 2004; Wesselingh, 2007). Another limitation of pollen supplementation studies is that they are often conducted on species with short flowering seasons or relatively few flowers. This is likely because comprehensive supplementation across a long season is extremely labour-intensive (Ashman et al., 2004). However, focusing on short-flowering species assumes that reproductive effort remains constant throughout the season. If pollen-supplemented flowers differ in seed set depending on the timing of pollination, it suggests that resource constraints play a greater role in determining reproductive output than pollen availability.

Seed provisioning may vary over the course of the flowering season due to changes in resource availability, pollen supply, or pollinator activity. In general, studies on natural populations have found that early-season flowers typically have greater seed set than late-season flowers (e.g., *Polemonium foliosissimum*; Zimmerman & Pyke, 1988). One hypothesis for this is that once plants are large enough to reproduce, they allocate heavily to reproduction, gradually depleting their resources as the season progresses (Medrano et al., 2000). Alternatively, early flowers may have greater access to resources due to their proximity to the basal, resource-rich part of the plant, whereas later, more distal flowers may receive fewer resources (Diggle, 1997). This pattern is most relevant in plants with acropetal flowering, where flowers develop sequentially from the base upward (Medrano et al., 2000). To separate the effects of pollen availability and resource limitation, experiments should track the timing of individual flower production and add supplemental pollen throughout an extended flowering season. Zimmerman and Pyke (1988) conducted four pollen supplementation treatments throughout the flowering season of *Polemonium foliosissimum* and found that both pollen-supplemented and control plants followed a similar pattern with the greatest seed set early in the season and declining toward the end. Together, this work suggests that declines in seed set across the season primarily reflect resource allocation within plants. However, to our knowledge no-one has investigated the effect of seasonal changes in seed provisioning within the framework of differences in plant resource status and life-history trade-offs.

Our primary goal in this study is to test whether plants that originate as seedlings versus clonal shoots (rhizomes) differ in their strategies of growth, survival, and reproduction, and whether these patterns differ within a growing season and across years. We predicted that seedlings would invest in faster growth and reproduction at the cost of overwinter survival, whereas clonal shoots would invest less in growth and reproduction initially and have greater survival. We also aimed to determine whether flower and seed production changed across the growing season, in natural open-pollinated plants and in the absence of pollen limitation, and to compare these patterns between seedlings and clonal shoots. We used the facultative clonal herb *Saponaria officinalis* (common soapwort), an invasive plant that produces underground rhizomes and flowers continuously through an extended season. To achieve our objectives, we grew plants that originated as seedlings or as rhizomes in common gardens and measured growth and reproduction over two years and survival over three years. We assigned a subset of plants to an open-pollination treatment and a subset to a supplemental-pollination treatment and recorded the date of opening for every flower and hand-pollinated nearly all open flowers in the supplemental-treatment to assess how flowering and seed production varied across the season and the implications on subsequent survival. Together, our results allow us to examine fitness components in plants that begin with very different resource status, providing insight into the factors shaping life-history strategies in clonal plants.

## Methods

### Study species

*Saponaria officinalis* L. (Caryophyllaceae) is a perennial herb native to Europe and parts of temperate Asia, and introduced to North America in the 1700s for its soap-making properties due to high saponin content (Mitich, 1990). It is now widespread and invasive across much of the continent. In both native and non-native habitats, it typically grows in sunny, open areas with well-drained soils, such as roadside ditches and field edges. In the region of Kingston, Ontario, plants emerge in early spring, with vegetative growth continuing until late fall. Flowering begins in mid-June and continues through to the first frost. Flowers are hermaphroditic and protandrous, beginning in male phase for 1–3 days, followed by female phase for another 2–3 days (Davis & Turner-Jones, 2008). Flowers are self-compatible and produce dry capsules containing 40–60 small seeds (1.5–2 mm in diameter), which are primarily gravity-dispersed. In addition to sexual reproduction, *S. officinalis* spreads clonally via underground rhizomes. The species often grows in dense patches, though the extent to which these represent clonal versus sexually derived individuals remains unknown (Lokker & Cavers, 1995).

### Sampling and plant measurements

In August 2021, we collected seed from 31 *S. officinalis* populations in the Kingston area (Table S1), selecting populations at least 500 m apart to reduce the likelihood of sampling from the same genetic population. Within each population, seeds were collected from plant stalks spaced at least one metre apart to increase the chance of sampling different genets. We obtained seed from 175 maternal plants and sowed seeds with the goal of growing four half-sibling replicates per individual. Seeds were stratified at 4 °C in moist sand for four weeks, then germinated in a growth chamber set to 22°C with a 16-hour photoperiod. After germination, seedlings were transferred to the Queen’s University Phytotron greenhouse and maintained under a 15.5-hour photoperiod at 21/18°C day/night temperatures. Following two weeks of growth, seedlings were vernalized for four weeks at 4 °C in the dark, then returned to the greenhouse for one week. During this final week, they were gradually hardened by increasing outdoor exposure from one hour on the first day to approximately eight hours by the end of the week. Only seedlings with at least two true leaves were transplanted.

In June 2022, we collected rhizome shoots from 25 populations, mostly overlapping with the seed collections. Within each population, we sampled from patches located at least one metre apart, treating each patch as a distinct individual (genet). From each genet, we collected four stalks (ramets) of similar size and temporarily potted them together in 4-inch pots for transport to the field site. In total, we collected 741 rhizome shoots from 185 maternal plants.

The experimental field site was located at Queen’s University Biological Station Bracken Tract meadow (44°38’36.8"N 76°20’06.5"W). The common garden was located within a large abandoned agricultural field in full sun. In May 2022, we mechanically tilled two 9.75 m × 9.75 m plots and covered them with ground cloth. We divided each plot in half with a 90 cm aisle for access and then subdivided each plot into 16 blocks (8 in each half). Using this split-plot design, each block was then alternatively assigned to contain plants that originated as seedlings or rhizomes. We cut square holes (10 cm × 10 cm) in the ground cloth spaced 60 cm apart, with each block containing 12 rows and four columns.

Planting occurred in mid-June 2022. We planted 699 seedlings from 175 maternal families (up to four half-sibling replicates per family) across 15 blocks. At the same time, we planted 742 rhizome shoots from 185 clonal individuals across 16 blocks. To reduce edge effects, we planted a border of extra individuals along the sides of the plots. Some rhizome shoots exhibited transplant stress and were temporarily supported with wooden skewers and twist ties. All individuals were watered daily for the first week after transplanting. In total, 1,441 individuals were planted in the common garden experiment.

One week after planting, we recorded initial size measurements for all individuals, including height, leaf count, and qualitative condition (e.g., signs of transplant stress). Transplant stress caused mortality in 157 plants (approximately equal numbers of seedlings and rhizomes) and were excluded from subsequent analyses. Throughout each growing season (June to September in 2022 and May to September in 2023), once a month we recorded height, relative size (on a 1–5 scale), the number of basal branches, deer herbivory level (on a 0–2 scale), and survival. Final survival was recorded in May 2024. For all 1,284 individuals in year one, we recorded whether each plant was reproductive (budding or flowering) approximately two weeks after the first flowers of the experiment opened. At the end of years one and two, we recorded whether plants flowered and categorized flowering intensity for those individuals as low, medium, or high relative to others within the same year. We randomly selected a subset of 55 individuals per growth type (seedling vs. rhizome shoot) for which we thoroughly examined reproductive output. In these individuals, we recorded final flower number, and counted the total number of fruit capsules, along with the number of filled versus empty capsules in both 2022 and 2023. However, some individuals died before they reproduced, resulting in a sample of 48 rhizomes and 53 seedlings.

### Pollen supplementation experiment

Within the larger common garden experiment, we conducted a comprehensive pollen supplementation experiment in 2022 to test i) whether plants are pollen limited; ii) the progression of flowering and seed set through the season in plants originating as seedlings versus rhizomes; and iii) whether there were costs of reproduction in the absence of pollen limitation. We randomly selected 60 rhizomes and 60 seedlings for this experiment, but a few individuals were subsequently damaged by deer and excluded. Half of the individuals within each growth type were assigned to receive supplemental pollen (29 rhizomes and 28 seedlings), while the other half remained open-pollinated (29 rhizomes and 29 seedlings). For supplemented plants, every flower received pollen from at least two non-half-sibling donor plants. To track flowering phenology, all flowers in both treatments were marked with a dot of nail polish on the calyx, with different colours representing the week of anthesis (e.g., week one flowers were marked red). The number of marked flowers was recorded weekly to monitor floral production and fruit retention. If a flower wilted before pollination could occur, it was marked as ‘missed’ for that week. We continued this procedure for five weeks, from August 8 to September 12, after which cool temperatures made it unlikely that later flowers would produce mature capsules.

When the fruit of *S. officinalis* is ripe, the top of the capsule splits open and drops seed. We aimed to harvest mature fruit prior to splitting. For each plant, we harvested seed from a given week together (i.e., capsules with the same nail polish colour were put into the same paper bag). We put ‘missed’ flowers from each week into different bags. Later, we sorted and counted filled or empty capsules. We then removed the seeds from filled capsules and removed debris with a small fan. We counted seed with an automated seed counter (Elmor C3 High Sensitive Seed Counter; Elmor Ltd.).

### Statistical analyses

Only individuals that survived beyond transplant stress were included in analyses (n = 1,284). Analyses were performed in R version 4.4.0 (R Core Team, 2024). Model fit was assessed during initial analyses and guided the choice of error distributions for each response variable. Continuous and log-transformed count responses were analyzed with linear mixed-effects models, and proportional responses with binomial generalized linear mixed-effects models. All mixed models were fit using the *lme4* package (Bates et al., 2015). Assumptions of homoscedasticity and residual normality in linear models, and dispersion in binomial models, were evaluated using the *DHARMa* package (Hartig, 2024). For all models, we extracted estimated marginal means and trends using the *emmeans* package (Lenth, 2025).

### Vegetative size

We assessed vegetative size by analyzing end-of-season height, branch number, and growth rate across the first two growing seasons (2022 and 2023). End-of-season height was analyzed with a linear mixed-effects model including growth type (seedling vs. rhizome), year, and their interaction as fixed effects, with plant ID and block as random effects. Branch number was log-transformed and analyzed with the same model structure. Growth rate was evaluated by modelling log-transformed height across time points, with linear and quadratic terms for time and their interactions with growth type and year, including plant ID and block as random effects.

### Reproduction

We first asked whether seedlings and rhizomes differed in their likelihood of reproducing early in the season. Reproductive status (reproductive or vegetative) was modeled using a binomial mixed-effects model with growth type as a fixed effect, and block and population as random effects. We then tested whether the size at which individuals initiated reproduction differed between growth types by comparing the log-transformed height of individuals that were reproductive using a linear mixed-effects model with the same fixed and random effects.

Our next set of analyses used the subset of ∼55 individuals of each growth type for which we counted fruit capsules. To test whether flower number differed between growth types and years, we modeled log-transformed capsule number with growth type, year, and their interaction as fixed effects, and plant ID as a random effect. We then calculated flowers per unit height (capsule number divided by plant height) and analyzed the log-transformed values using the same model structure. Finally, we compared the proportion of fruit capsules filled using a binomial generalized linear mixed-effects model with growth type, year, their interaction, and standardized height as fixed effects, and plant ID as a random effect.

We assessed pollen limitation and seasonal variation in reproductive output using the subset of 60 ramets and 60 seedlings, with 30 open-pollinated and 30 pollen-supplemented individuals per growth type. We examined four measures of reproductive output for each week of flower opening, calculated from the weekly harvests of capsules (i.e., non-cumulative values): (1) flower number, measured as the total number of fruit capsules; (2) the proportion of capsules filled with seeds; (3) average seed set per pod, calculated as the number of seeds divided by the number of filled capsules; and (4) total seed number. All models included growth type, pollination treatment, week, and their interactions, with relative size (scored from 1–5) as a covariate. A quadratic term for week was tested in each model and retained only if significant. Plant ID and block were included as random effects. Log-transformed flower number, seed set per pod, and log-transformed seed number were analyzed using linear mixed-effects models, while fruit set was analyzed using a binomial mixed-effects model. In addition to weekly measures, we also analyzed total reproductive output across the season by summing seeds per plant across weeks. Total seed number was log-transformed and fit with a linear model including growth type, pollination treatment, and their interaction, with relative size as a covariate.

### Survival

We assessed differences in survival between rhizomes and seedlings that survived beyond transplant stress (n = 1,284) across three years. Survival was recorded at five key timepoints: one month after planting, the end of the first growing season, overwinter survival into year two, the end of the second growing season, and overwinter survival into year three. We first modeled survival of all individuals using a Cox proportional hazards model (*coxme* package; Therneau, 2024) with growth type as the main predictor and block as a random effect. To evaluate whether additional traits influenced survival beyond growth type, we then ran a second Cox model restricted to the individuals that survived past the first timepoint (n = 1,270). In this model, fixed effects included growth type and end-of-year-one traits: plant height, basal branch number, flowering intensity category (none, low, medium, or high), and herbivory, with block included as a random effect. Finally, we ran a third model including only individuals in the pollen-supplementation experiment (n = 116) to test whether manipulated reproductive effort influenced survival. Model structure was similar to the second model, but we added pollination treatment and, instead of flowering intensity category, included the total number of flowers and seeds as predictors. Block was retained as a random effect.

## Results

### Vegetative Size

By the end of the first year, seedlings and rhizomes did not differ significantly in height, but seedlings had more branches than rhizomes (Figure 1A; Table 1; Table S2). Seedlings also grew faster than rhizomes during the first year (Figure 1A; Table 1; Table S2). Comparisons between years showed both growth types increased significantly in height and growth rate in their second year, but while rhizomes also increased in branch number, seedlings made fewer branches in year two than in year one (Table S3). By the end of the second year, rhizomes were taller, had more branches, and had grown faster than seedlings (Figure 1; Table 1; Table S2).

**Figure 1.**
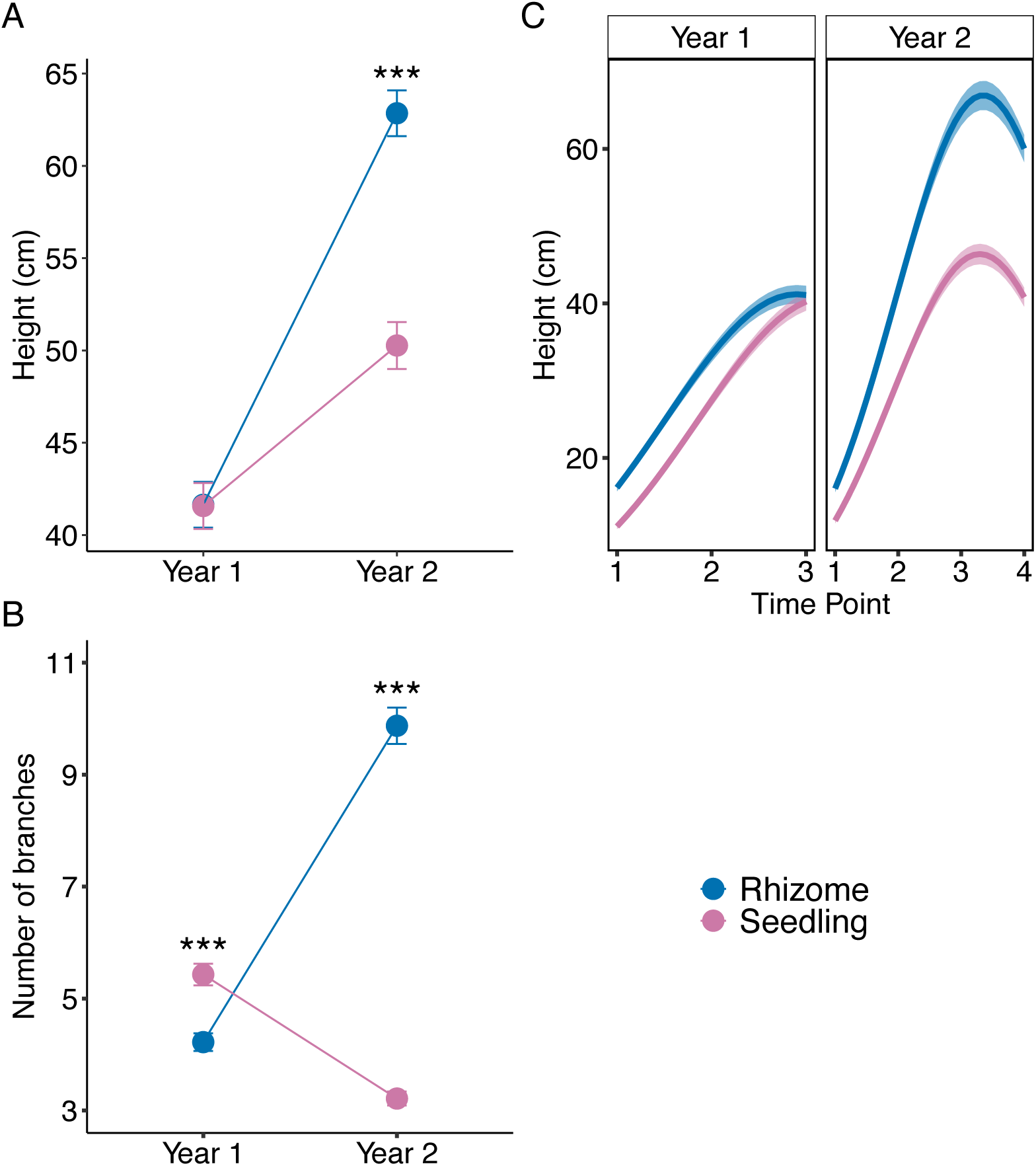
Differences in (A) height, (B) branch number, and (C) height trajectories through time, between rhizomes and seedlings over two growing seasons. Points show estimated marginal means with standard errors and curves in panel C indicate estimated slopes with standard error ribbons. Branch number and height trajectories are back-transformed from model predictions. Rhizomes are shown in blue, and seedlings in pink. Asterisks indicate significance levels for pairwise contrasts between growth types within years: *p* < 0.05*, *p* < 0.01**, *p* < 0.001***. See Table 1 for statistical details.

**Table 1.** Linear mixed-models (*F*-statistics with *p*-values) for vegetative traits of 1,284 *Saponaria officinalis* grown in a common garden. Statistically significant *p*-values are shown in bold. See Figure 1.

| Model | df | $F$-value | $p$ |
| --- | --- | --- | --- |
| **1. Height** |  |  |  |
| Growth type | 1, 1228.3 | 132.23 | **<0.001** |
| Year | 1, 1176.9 | 1242.49 | **<0.001** |
| Growth type × Year | 1, 1176.8 | 217.60 | **<0.001** |
| **2. Branch number** |  |  |  |
| Growth type | 1, 849.6 | 145.90 | **<0.001** |
| Year | 1, 1168.2 | 37.57 | **<0.001** |
| Growth type × Year | 1, 1168.2 | 519.56 | **<0.001** |
| **3. Growth rate** |  |  |  |
| Growth type | 1 4814.1 | 371.80 | **<0.001** |
| Year | 1 6798.0 | 28.73 | **<0.001** |
| Time point | 1 6783.1 | 7330.55 | **<0.001** |
| Time point<sup>2</sup> | 1 6782.8 | 3763.29 | **<0.001** |
| Growth type × Year | 1 6799.7 | 64.48 | **<0.001** |
| Growth type × Time point | 1 6784.2 | 98.76 | **<0.001** |
| Year × Time point | 1 6777.1 | 232.15 | **<0.001** |
| Growth type × Year × Time point | 1 6781.4 | 192.63 | **<0.001** |

### Reproduction

Seedlings and rhizomes did not differ in the probability of being reproductive early in the first growing season (χ^2^ = 0.0034, *p* = 0.95). However, among reproductive individuals, seedlings were significantly shorter at budding than rhizomes (*F*_1,329.9_ = 43.789, *p* < 0.001). By the end of the first year 91.1% of surviving rhizomes and 88.4% seedlings had flowered.

In the random subset of individuals for which we more thoroughly assessed reproductive output, we found that seedlings had significantly more flowers and flowers per unit height than rhizomes in the first growing season (Figure 2; Table 2; Table S2). However, rhizomes had a higher proportion of capsules filled than seedlings in the same year. By the second year, rhizomes had significantly greater flower number and flowers per unit height, whereas seedlings did not differ across years (Figure 2; Table S3). In the second growing season, rhizomes had more flowers than seedlings, but the two growth types did not differ in flowers per unit height or in the proportion of capsules filled (Figure 2; Table 2; Table S2). Neither growth type showed a significant change in the proportion of capsules filled between years (Table S3).

**Figure 2.**
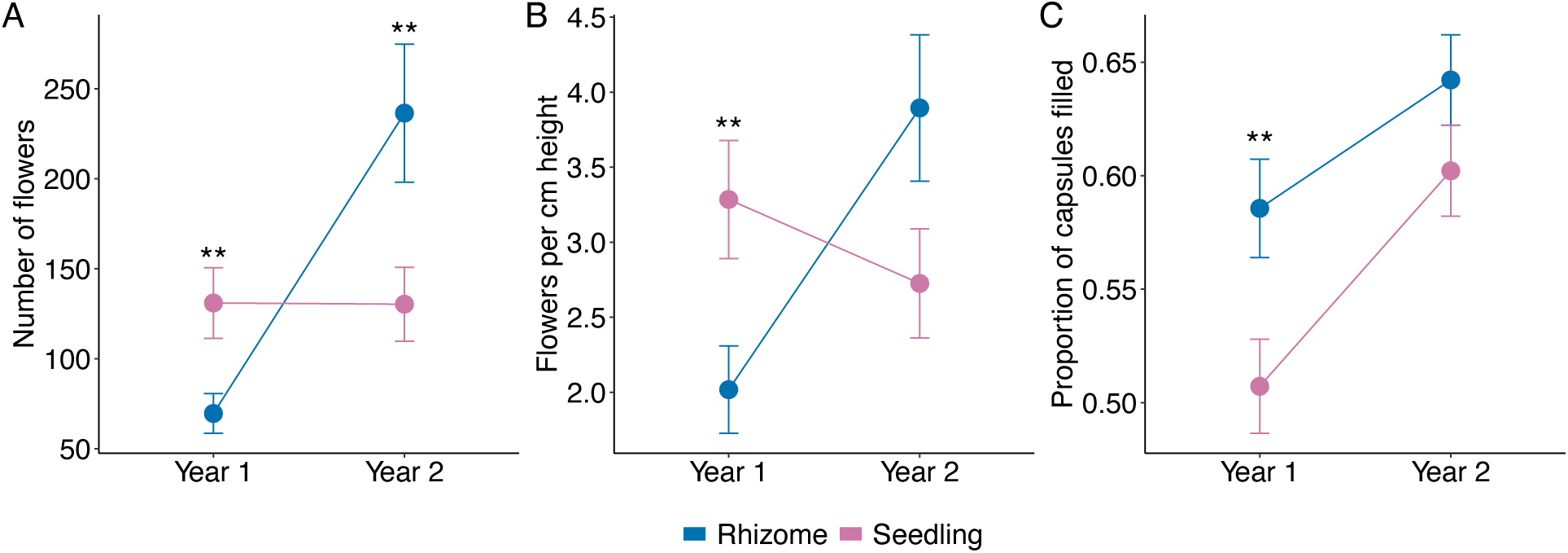
Differences in (A) the number of flowers, (B) the number of flowers per cm of height and (C) the proportion of capsules filled, in rhizomes and seedlings across two growing seasons. Points represent estimated marginal means with standard errors. Flower number and flowers per cm of height are back-transformed from model predictions. Rhizomes are shown in blue, and seedlings in pink. Asterisks indicate significance levels for pairwise contrasts between growth types within years: *p* < 0.05*, *p* < 0.01**, *p* < 0.001***. See Table 2 for statistical details.

**Table 2.**
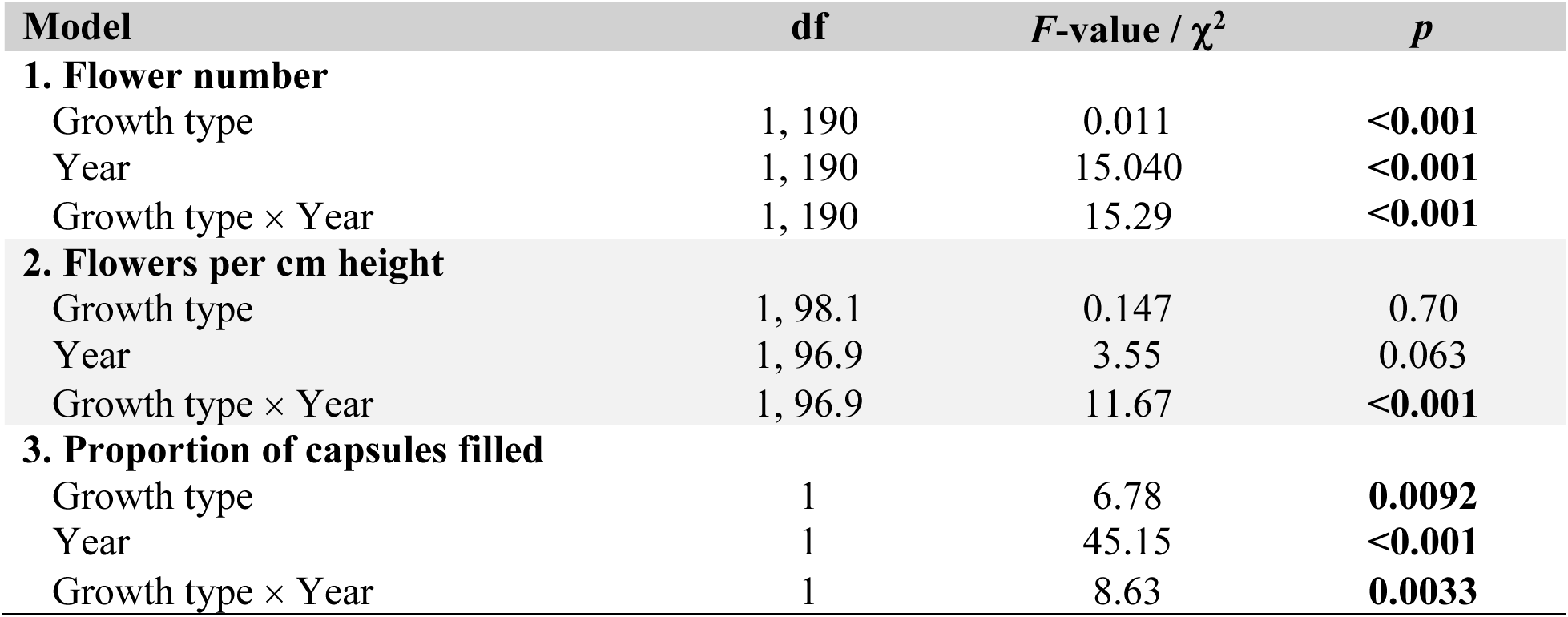
Linear and generalized linear mixed-models (*F*- and χ^2^-tests with *p*-values) for reproductive traits of a subset of 101 *Saponaria officinalis* individuals. Statistically significant *p*-values are shown in bold. See Figure 2.

| Model | df | $F$-value / $\chi^2$ | $p$ |
| --- | --- | --- | --- |
| **1. Flower number** |  |  |  |
| Growth type | 1, 190 | 0.011 | **<0.001** |
| Year | 1, 190 | 15.040 | **<0.001** |
| Growth type × Year | 1, 190 | 15.29 | **<0.001** |
| **2. Flowers per cm height** |  |  |  |
| Growth type | 1, 98.1 | 0.147 | 0.70 |
| Year | 1, 96.9 | 3.55 | 0.063 |
| Growth type × Year | 1, 96.9 | 11.67 | **<0.001** |
| **3. Proportion of capsules filled** |  |  |  |
| Growth type | 1 | 6.78 | **0.0092** |
| Year | 1 | 45.15 | **<0.001** |
| Growth type × Year | 1 | 8.63 | **0.0033** |

In the pollen supplementation experiment, all four reproductive traits were positively associated with plant size, and each trait varied significantly across the season (linear or quadratic week effects; Table 3). Flower number increased across the season but did not differ between growth types or pollination treatments (Figure 3A; Table 3). The proportion of capsules filled varied significantly between pollination treatments and growth types across the season (Figure 3B; Table 3). Pollen-supplemented seedlings consistently had the greatest proportion of capsules filled the first three weeks, exceeding all other growth type × treatment combinations (pairwise contrasts, *p* < 0.05). In contrast, pollen-supplemented rhizomes showed a lower proportion of capsules filled relative to all other growth type × treatment combinations during the first three weeks. Open-pollinated seedlings and rhizomes did not differ from each other until the final week when seedlings had a greater proportion of capsules filled (*p* = 0.022). Average seed set per filled capsule declined significantly across the season for all growth type × treatment groups (Figure 3C; Table 3). Pollen-supplemented seedlings had greater average seed set per filled capsule than all other growth type × treatment combinations during the first three weeks (pairwise contrasts, *p* < 0.05). Total seeds increased across the season for all growth type × treatment combinations (Figure 3D; Table 3). In weeks four and five open-pollinated rhizomes made fewer seeds than open-pollinated seedlings (pairwise contrasts, *p* < 0.05). Total reproductive output, measured as the total number of seeds per plant, differed between growth × treatment combinations (Table S4). Open-pollinated rhizomes matured fewer seeds than open-pollinated seedlings (*p* = 0.0024) and pollen-supplemented rhizomes (*p* = 0.020). In contrast, open-pollinated seedlings matured more seeds than pollen-supplemented seedlings (*p* = 0.032).

**Figure 3.**
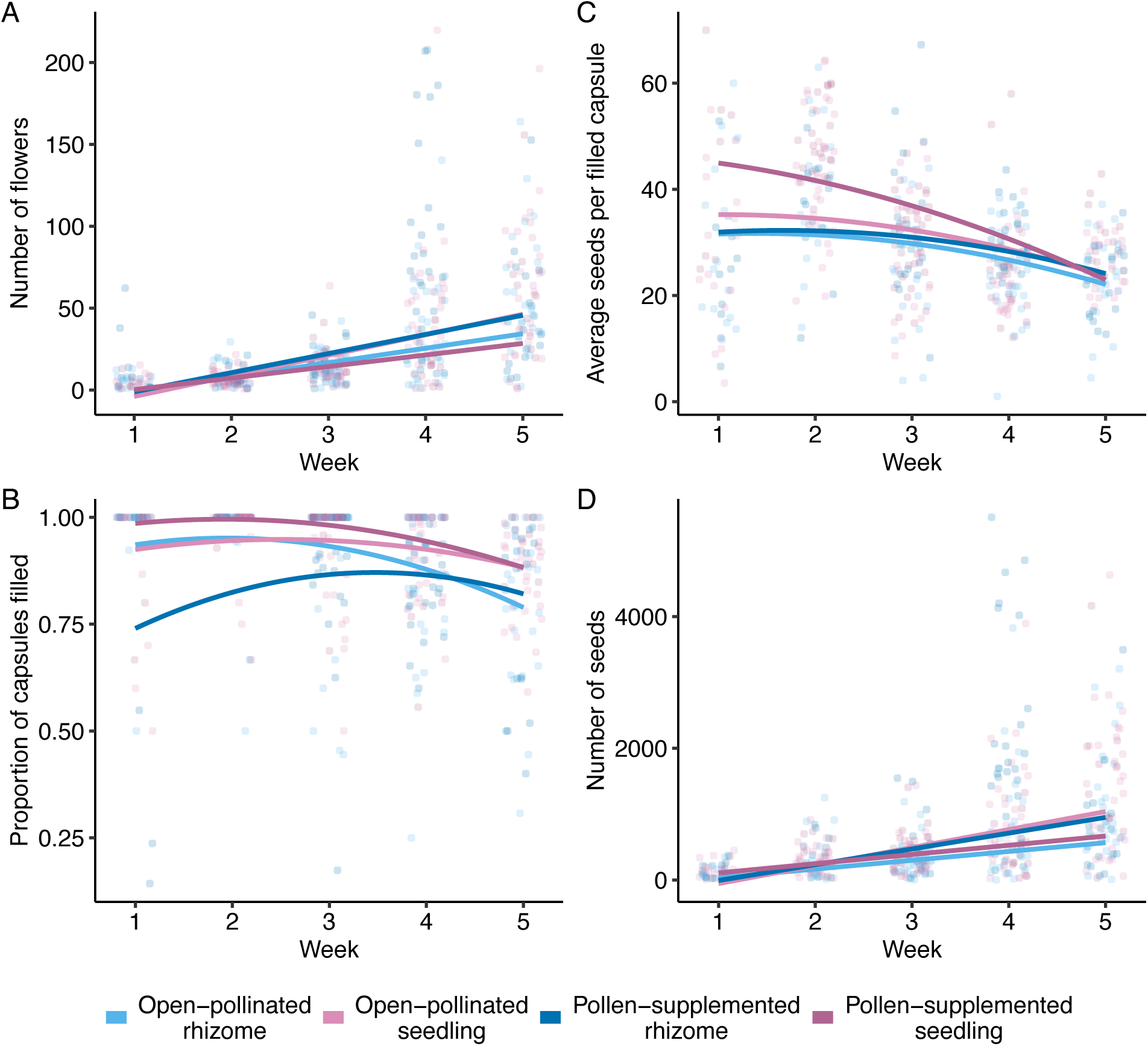
Results from the pollination experiment with rhizomes and seedlings that were open-pollinated or received supplemental pollen. Figures show values for each week of (A) number of flowers, (B) proportion of capsules filled with seed, (C) average seeds per filled capsule, and (D) total seed number. Points show individual-level raw data; lines represent fitted linear or quadratic trends based on model predictions at each week. Flower number and seed number are back-transformed from model predictions. Rhizomes are shown in blue, seedlings are in pink; darker colours are pollen-supplemented, lighter colours are open-pollinated. See Table 3 for statistical details.

**Table 3.** Linear and generalized linear mixed-models (*F*- and χ^2^-tests with *p*-values) for reproductive traits of 115 *Saponaria officinalis* in the pollen supplementation experiment. Statistically significant *p*-values are shown in bold. See Figure 3.

| Model | df | $F$-value / $\chi^2$ | $p$ |
| --- | --- | --- | --- |
| **1. Flower number** |  |  |  |
| Growth type | 1, 406.4 | 0.64 | 0.43 |
| Treatment | 1, 406.7 | 1.48 | 0.22 |
| Week | 1, 333.2 | 356.58 | **<0.001** |
| Size score (1-5) | 1, 109.2 | 10.13 | **0.0019** |
| Growth type × Treatment | 1, 406.8 | 0.0004 | 0.98 |
| Growth type × Week | 1, 333.1 | 0.10 | 0.75 |
| Treatment × Week | 1, 333.3 | 1.63 | 0.20 |
| Growth type × Treatment × Week | 1, 332.0 | 1.75 | 0.19 |
| **2. Proportion of capsules filled** |  |  |  |
| Growth type | 1 | 1.094 | 0.30 |
| Treatment | 1 | 27.73 | **<0.001** |
| Week | 1 | 7.174 | **0.0074** |
| Week<sup>2</sup> | 1 | 35.35 | **<0.001** |
| Size score (1-5) | 1 | 4.27 | **0.039** |
| Growth type × Treatment | 1 | 29.74 | **<0.001** |
| Growth type × Week | 1 | 7.67 | **0.0056** |
| Treatment × Week | 1 | 35.35 | **<0.001** |
| Growth type × Treatment × Week | 1 | 28.14 | **<0.001** |
| **3. Seed set per filled capsule** |  |  |  |
| Growth type | 1, 405.9 | 11.97 | **<0.001** |
| Treatment | 1, 405.9 | 4.17 | **0.042** |
| Week | 1, 38.7 | 0.33 | 0.57 |
| Week<sup>2</sup> | 1, 553.9 | 4.66 | **0.032** |
| Size score (1-5) | 1, 585.1 | 4.92 | **0.029** |
| Growth type × Treatment | 1, 502.3 | 4.23 | **0.040** |
| Growth type × Week | 1, 672.8 | 5.66 | **0.018** |
| Treatment × Week | 1, 187.5 | 1.58 | 0.21 |
| Growth type × Treatment × Week | 1, 358.5 | 3.02 | 0.083 |
| **4. Seed number** |  |  |  |
| Growth type | 1, 406.1 | 0.5417 | 0.46 |
| Treatment | 1, 406.4 | 2.7315 | 0.099 |
| Week | 1, 326.0 | 186.7628 | **<0.001** |
| Size score (1-5) | 1, 101.7 | 15.7940 | **0.0013** |
| Growth type × Treatment | 1, 406.6 | 1.5775 | 0.21 |
| Growth type × Week | 1, 325.9 | 0.0489 | 0.82 |
| Treatment × Week | 1, 326.0 | 1.3647 | 0.24 |
| Growth type × Treatment × Week | 1, 324.7 | 5.2901 | **0.022** |

### Survival

Seedlings had markedly lower survival than rhizomes: of the individuals that survived past transplant stress, approximately 27% of seedlings and 3% of rhizomes died over the course of the experiment. In the global Cox model including all individuals, seedlings were over seven times more likely to die at any given time than rhizomes (HR = 7.13, 95% CI = 4.96 – 10.25, *p* < 0.001; Figure 4). When plant traits measured at the end of the first season were included, seedlings again showed much lower survival (HR = 9.58, 95% CI = 6.34 – 14.47, *p* < 0.001). Taller plants and those with more basal branches had significantly greater survival (HR per cm = 0.95, 95% CI = 0.93 – 0.97, *p* < 0.001; HR per branch = 0.92, 95% CI = 0.86 – 0.99, *p* = 0.035), while flowering intensity and herbivory were not significant predictors. Finally, in the pollen-supplementation experiment, seedlings again had lower survival than rhizomes (HR = 32.0, 95% CI = 3.68 – 278.80, *p* = 0.002), and taller plants were more likely to survive (HR per cm = 0.86, 95% CI = 0.78 – 0.94, *p* = 0.002). Survival was not significantly influenced by pollination treatment or by the number of flowers, seeds, or branches in this subset. Although survival was not influenced by pollination treatment overall, pollen-supplemented seedlings were the only group to experience overwinter mortality between year one and year two (over 10% vs. 0% in the other groups).

**Figure 4.**
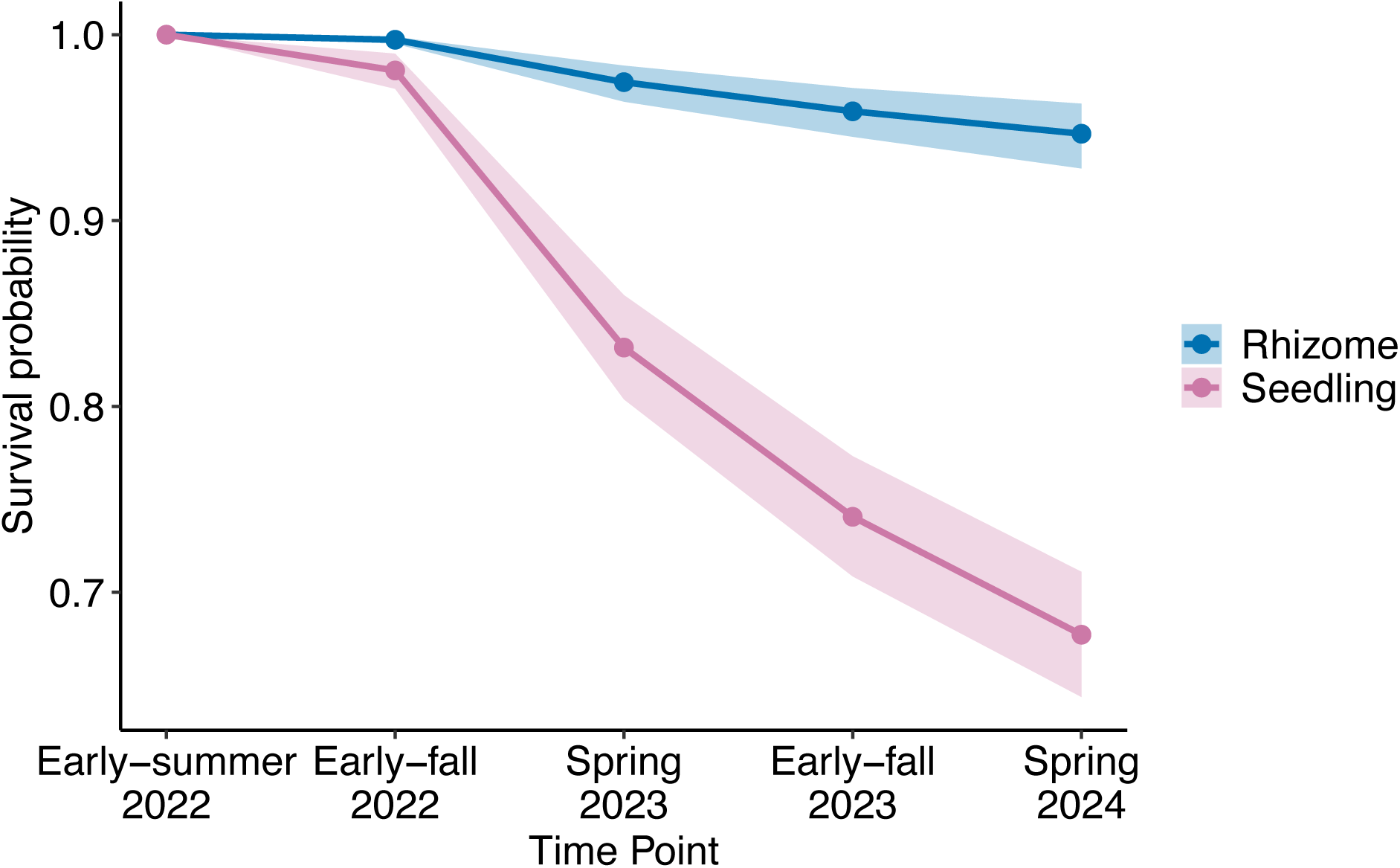
Survival analysis of rhizome ramets and seedlings across three years. Points show model-estimated survival probabilities at five time points, with ribbons indicating 95% confidence intervals based on 1,000 bootstrap replicates.

## Discussion

Our results are consistent with the hypothesis that growth origin influences subsequent life history strategy in *Saponaria officinalis*, with rhizomes and seedlings differing in growth, survival, and reproduction, within and across seasons. In the first year, individuals that originated as seedlings grew faster, initiated reproduction at a shorter height and made more flowers than rhizomes, indicative of an allocation strategy favouring early reproduction. Aligning with these patterns, seedlings had poorer establishment with lower survival, particularly overwinter between year one and two. For both growth types, end-of-season height was a significant predictor of overwinter survival, while flowering intensity and herbivory had no significant effect. These findings suggest that allocation to vegetative growth buffered plants against winter mortality.

In the subset of individuals used in the pollination experiment, average seed set per filled capsule declined over the flowering season for both growth types and treatments, suggesting that resource limitation played a larger role in restricting seed set than pollen limitation. Pollen-supplemented seedlings had greater seed set early in the season than open-pollinated seedlings or rhizomes, indicating that seedlings capitalize on favourable early pollination conditions. Notably, pollen-supplemented seedlings were the only group to experience overwinter mortality between year one and two, suggesting that increased reproductive effort came at a cost to survival.

### Growth origin affects life history strategies

We found that seedlings grew faster and bolted at a shorter height than rhizomes, in the first year of the experiment. For seedlings of clonal species, rapid growth and early reproduction may be essential for securing establishment, because seedlings have to compete with the parent plant and nearby seedlings and clonal shoots when dispersal distances are short (Qi et al. 2014). Even for species with farther dispersal, rapid early growth by seedlings may be favoured to deal with environmental unpredictability (Gross, 1984; Fenner, 1987). Rhizomes, by contrast, are supported by resources from the parent plant, making establishment more reliable (Lei, 2010). Across clonal perennials, recruitment by seedlings in established populations is generally rare and is often associated with disturbance (Harper, 1977; Eriksson, 1989). For example, seedlings were more abundant in grazed sites than in undisturbed alpine grasslands (Wu et al., 2011), and computer simulations suggest that high densities of *Ranunculus repens* genets occur only immediately after large-scale disturbance (Watkinson & Powell, 1993). These patterns suggest that *S. officinalis* seedlings in our experiment benefitted from the resource-rich, disturbed habitat created by tilling the garden before planting. Seedlings behaved in a manner consistent with Grime’s (1977) model of ruderal strategies, in which species in disturbance-prone habitats grow quickly and reproduce early.

Stress is another factor that can influence reproductive timing. In perennial plants, moderate stress often delays flowering (Bazzaz et al., 1987; Bennett et al., 2012), whereas extreme stress or low probability of survival can trigger a “reproductive burst” (Bazzaz et al., 1987). In our experiment, seedlings produced more flowers per unit height than rhizomes in the first year, indicating greater relative allocation to reproduction. This greater relative allocation is consistent with a strategy to maximize fitness when future survival is uncertain. However, despite larger investment in flowers, seedlings filled a smaller proportion of pods than rhizomes. This discrepancy suggests that limited resources prevented maturation of all ovules into seeds. It may also reflect stress-induced abortion of developing fruits, a response documented in both crops and wild species (Sparks, 1989; Sun et al., 2004).

Further support that seedlings face greater stress is that they experienced lower survival than rhizomes across the experiment. Note that we germinated seedlings in the greenhouse so that this does not include the most vulnerable phase of seedling development (Fenner, 1985). Similar patterns of lower seedling survival compared with established clonal shoots have been reported in *Salix exigua* (Douhovnikoff et al., 2005), *Wedelia trilobata* (Qi et al., 2008), and *Trifolium repens* (Turkington et al., 1979) and is consistent with life-history theory predicting low seedling survival in long-lived species (Deevey, 1947). However, variation in survival probability within growth types suggests that other factors are also at play.

Variation in resource acquisition can mask underlying trade-offs between growth, reproduction, and survival (Van Noordwijk & de Jong, 1986). In *Orchis purpurea*, for example, plants in high-light environments flowered at a smaller size, and flowering was associated with increased survival, whereas in low-light environments reproduction came with a survival cost (Jacquemyn et al., 2010). Consistent with these patterns, we found that taller, more branching individuals of *S. officinalis* were more likely to survive, regardless of whether they originated from seeds or rhizomes. In seedlings, some of the variation in vigour may reflect maternal effects through seed provisioning, which has been widely shown to influence seedling vigour (Wulff, 1986; Eriksson, 1999; Paz & Martinez-Ramos, 2003). In rhizomes, initial size at planting likely contributed to end-of-season variation despite efforts to standardize size. For example, larger rhizome fragments of *Clintonia borealis* had significantly higher survival than smaller ones (Ashmun & Pitelka, 1985). In *S. officinalis*, future resource manipulation experiments could help determine whether the links between height, branching and survival are due to differences in allocation strategies or reflect access to greater resource pools. Still, the shift from seedlings exhibiting greater vegetative and reproductive traits in year one to rhizomes surpassing them in year two suggests that allocation differences are likely involved.

By the second year of the experiment, rhizomes grew faster, were taller and had more branches and flowers than seedlings. This difference between rhizomes and seedlings may reflect greater belowground investment by rhizomes, fuelling subsequent growth. Evidence from other species supports this idea: in *Aster acuminatus* and *Clintonia borealis*, genet size increased from year to year when grown from rhizome fragments (Ashmun & Pitelka, 1984, 1985). Similarly, in *Agropyron repens* and *A. caninum*, plants established from rhizomes allocated a greater proportion of total biomass to belowground structures than those established from seed (Tripathi & Harper, 1973). Although we did not measure belowground biomass in *S. officinalis*, the greater year-two performance of rhizomes is consistent with conservative resource allocation in the first year, followed by using stored reserves for overwinter survival and subsequent growth (Obeso, 2002). Future work comparing belowground biomass within and between growth types could help clarify the mechanisms driving differences in growth and survival in *S. officinalis*.

### Costs of reproduction within and between seasons

Seasonal declines in provisioning flowers are common in angiosperms, with later flowers typically smaller and producing fewer ovules, seeds, and fruits than earlier ones (Ashman & Baker, 1992; Diggle, 1995). Consistent with this pattern, we found that for both rhizomes and seedlings of *S. officinalis* average seed number per capsule decreased over time. The decline is unlikely to be caused by insufficient pollination, because both open pollinated and supplementally pollination plants displayed the same pattern. This suggests that it is due to resource constraints on reproduction. Position-dependent effects may contribute, as later flowers in *S. officinalis* develop farther from basal sources of resource supply (Diggle, 1995; Kliber & Eckert, 2004). At the same time, competition may intensify among simultaneously developing flowers later in the season, as shown by the decline in seed number per capsule alongside increasing flower number. A previous experiment in this species that removed flowers (to simulate herbivore damage) found greater seed mass when flowers were removed (Lokker & Cavers, 1995). Together, these findings suggest that fewer seeds per capsule reflects both sequential competition between early and late flowers and simultaneous competition among developing fruits.

An additional factor that could influence seasonal patterns of seed set in *S. officinalis* is its protandrous flowers. Protandry is a form of dichogamy in which hermaphroditic flowers present male function first and then transition to female phase. In protandrous species, sex ratios typically shift from male- to female-dominated over the flowering season, a dynamic that can induce negative frequency-dependent selection favouring early flowers that preferentially invest in female function (Brunet & Charlesworth, 1995; Ishii & Harder, 2012). In our experiment, greater seed number per capsule early in the season is consistent with higher investment in female function at that time. A next step would be to examine seed mass from this experiment to determine whether early-season seeds were better provisioned.

Life-history theory predicts that current reproductive investment can impose costs on future survival or fecundity (Williams, 1966; Stearns, 1989). Costs have been found in many studies (see Dorken et al., 2025) but are more likely to be detected under specific environmental conditions (Sandvik, 2001; Sletvold & Ågren, 2015; Hamann et al., 2021) or when reproductive effort is experimentally manipulated (Primack & Hall, 1990; Horvitz et al., 2010). In *S. officinalis*, overall survival did not differ significantly between pollination treatments. However, pollen-supplemented seedlings were the only group to experience overwinter mortality between the first and second growing seasons, despite showing no obvious differences in size or reproductive output relative to other individuals. This pattern suggests that while early investment by pollen-supplemented seedlings may have boosted short-term reproductive returns, it also carried hidden costs.

### Conclusions

This study highlights the value of comparing seedlings and clonal shoots to understand the life history strategies of invasive perennials. By tracking growth, reproduction and survival across multiple years, we show how *S. officinalis* may combine establishment from seed with long-term clonal persistence. However, whether *S. officinalis* populations are maintained by ongoing seed dispersal or are largely the product of initial founding events followed by clonal spread remains unknown. Genetic analysis across populations could clarify the number of founding genets, patterns of clonal dominance, and the relative roles of sexual and asexual reproduction in demographic dynamics. Overall, these findings highlight the importance of early establishment conditions and the potential for sexual and asexual reproductive strategies to support persistence and spread. This approach has broad relevance for studying population dynamics and can inform both management strategies and restoration efforts.

## Author Contributions

A.V.N. and J.F. conceived the project; A.V.N. collected the data; A.V.N. analyzed the data; A.V.N. and J.F. wrote the manuscript; J.F. oversaw all aspects of the project. Both authors approved the final version of the manuscript.

## Acknowledgements

The authors thank I. Lewis, J. Lee, E. Gillete, A. Pagulayan, B. Balcaran, C. Ma, S. Gama, B. Badali, R. Marcelissen, and R. Cross for help collecting data; and the Queen’s University Biological Station for access to the field and logistical support. The authors also thank colleagues C. Eckert, S. Yakimowski for helpful suggestions throughout the project.

## Supplementary Material

**Table S1.** Populations of *Saponaria officinalis* used in the common garden experiment and their geographical coordinates. Population letters refer to the sampling region (B- Battersea, E- Elbow Lake, P- Opinicon Road, S- Syndeham). Approximately four individuals were collected from each maternal family for both rhizomes and seedlings.

| Population | Number of maternal families - rhizomes | Number of maternal families - seedlings | Latitude (°N) | Longitude (°W) |
| --- | --- | --- | --- | --- |
| B1 | 12 | 6 | 44.286709 | 76.465904 |
| B2 | 2 | 0 | 44.313029 | 76.464727 |
| B3 | 17 | 2 | 44.346021 | 76.441621 |
| B4 | 2 | 5 | 44.341779 | 76.442625 |
| B5 | 3 | 2 | 44.402466 | 76.408243 |
| B6 | 1 | 1 | 44.423825 | 76.372993 |
| B7 | 9 | 10 | 44.430473 | 76.373520 |
| B8 | 0 | 18 | 44.405587 | 76.408421 |
| B9 | 3 | 1 | 44.378191 | 76.429067 |
| B10 | 2 | 1 | 44.356047 | 76.430254 |
| B11 | 2 | 6 | 44.352613 | 76.424414 |
| B12 | 0 | 2 | 44.268048 | 76.479555 |
| E1 | 0 | 1 | 44.428716 | 76.465769 |
| E2 | 0 | 1 | 44.429058 | 76.465110 |
| E3 | 0 | 1 | 44.432315 | 76.459979 |
| E4 | 9 | 5 | 44.429132 | 76.465081 |
| E5 | 2 | 0 | 44.432303 | 76.460154 |
| E6 | 13 | 9 | 44.457882 | 76.428570 |
| P1 | 3 | 3 | 44.481493 | 76.478884 |
| P2 | 0 | 2 | 44.483267 | 76.473476 |
| P3 | 16 | 17 | 44.484425 | 76.472469 |
| P4 | 0 | 4 | 44.484080 | 76.455651 |
| P5 | 0 | 1 | 44.486581 | 76.449537 |
| P6 | 3 | 5 | 44.488808 | 76.441085 |
| S1 | 9 | 8 | 44.407035 | 76.573157 |
| S2 | 17 | 10 | 44.407533 | 76.591162 |
| S3 | 11 | 6 | 44.475382 | 76.561532 |
| S4 | 6 | 0 | 44.518016 | 76.579484 |
| S5 | 11 | 8 | 44.521426 | 76.582032 |
| S6 | 2 | 9 | 44.406359 | 76.61637 |
| S7 | 10 | 11 | 44.407364 | 76.56525 |
| S8 | 13 | 11 | 44.412806 | 76.527532 |
| S9 | 0 | 1 | 44.424908 | 76.527363 |
| S10 | 10 | 8 | 44.450826 | 76.508371 |
| S11 | 0 | 1 | 44.389500 | 76.572600 |

**Table S2.** Pairwise contrasts between seedlings and rhizomes within each year. Contrasts are based on estimated marginal means from mixed-effects models of vegetative and reproductive traits (See Table and Table). Vegetative traits include all plants, while reproductive traits were assessed in a random subset of ∼55 individuals of each growth type. Positive *t*/*z* -values indicate greater trait values in seedlings, and negative values indicate greater trait values in rhizomes. Statistically significant *p*-values are shown in bold.

| **Trait** | **Year** | **df** | **$t/z$-ratio** | **$p$-value** |
| --- | --- | --- | --- | --- |
| **Vegetative** |  |  |  |  |
| Height | 2022 | 2157 | 0.098 | 0.92 |
| Height | 2023 | 2233 | 17.40 | **<0.001** |
| Number of branches | 2022 | 1916 | -5.16 | **<0.001** |
| Number of branches | 2023 | 1863 | 23.79 | **<0.001** |
| Growth rate | 2022 | 6922 | -14.43 | **<0.001** |
| Growth rate | 2023 | 6940 | 3.49 | **<0.001** |
| **Reproductive** |  |  |  |  |
| Flower number | 2022 | 190 | -2.90 | **0.0042** |
| Flower number | 2023 | 190 | 2.64 | **0.0091** |
| Flowers per unit height | 2022 | 187 | -2.63 | **0.0092** |
| Flowers per unit height | 2023 | 188 | 1.96 | 0.052 |
| Proportion of capsules filled | 2022 | - | 2.60 | **0.0092** |
| Proportion of capsules filled | 2023 | - | 1.42 | 0.16 |

**Table S3.** Pairwise contrasts within growth types between years one and two. Contrasts are based on estimated marginal means from mixed-effects models of vegetative and reproductive traits (see Table and Table). Vegetative traits include all plants, while reproductive traits were assessed in a random subset of ∼55 individuals of each growth type. Positive *t*/*z* -values indicate greater trait values in year one, and negative values indicate greater trait values in year two. Statistically significant *p*-values are shown in bold.

| **Trait** | **Growth type** | **df** | ***t/z*-ratio** | ***p*-value** |
| --- | --- | --- | --- | --- |
| **Vegetative** |  |  |  |  |
| Height | Rhizomes | 1099 | -37.108 | **<0.001** |
| Height | Seedlings | 1262 | -13.865 | **<0.001** |
| Number of branches | Rhizomes | 1265 | -20.387 | **<0.001** |
| Number of branches | Seedlings | 1274 | 11.821 | **<0.001** |
| Growth rate | Rhizomes | 6910 | -20.892 | **<0.001** |
| Growth rate | Seedlings | 6938 | -2.358 | **0.018** |
| **Reproduction** |  |  |  |  |
| Flower number | Rhizomes | 96.1 | -5.39 | **<0.001** |
| Flower number | Seedlings | 97.8 | 0.023 | 0.98 |
| Flowers per unit height | Rhizomes | 94.8 | -3.69 | **<0.001** |
| Flowers per unit height | Seedlings | 97.4 | 1.10 | 0.27 |
| Proportion of capsules filled | Rhizomes | - | -6.720 | **<0.001** |
| Proportion of capsules filled | Seedlings | - | -11.022 | **<0.001** |

**Table S4.** Pairwise contrasts of total seed production across growth type × pollination treatment combinations from the pollen-supplementation experiment. Contrasts are based on estimated marginal means from a mixed-effects model that includes relative size as a covariate. Statistically significant *p*-values are shown in bold.

| **Contrast** | **df** | ***t*-ratio** | ***p*-value** |
| --- | --- | --- | --- |
| Open-pollinated rhizome – Open-pollinated seedling | 110 | -3.63 | **0.0024** |
| Open-pollinated rhizome – Pollen-supplemented rhizome | 110 | -2.95 | **0.019** |
| Open-pollinated rhizome – Pollen-supplemented seedling | 110 | -0.82 | 0.85 |
| Open-pollinated seedling – Pollen-supplemented rhizome | 110 | 0.67 | 0.91 |
| Open-pollinated seedling – Pollen-supplemented seedling | 110 | 2.78 | **0.032** |
| Pollen-supplemented rhizome – Pollen-supplemented seedling | 110 | 2.085 | 0.16 |

## References

Aragón, C. F., Méndez, M., & Escudero, A. (2009). Survival costs of reproduction in a short-lived perennial plant: live hard, die young. American Journal of Botany, 96, 904–911.

Ashman, T.-L. (1994). A dynamic perspective on the physiological cost of reproduction in plants. American Naturalist, 144, 300–316.

Ashman, T.-L., & Baker, I. (1992). Variation in floral sex allocation with time of season and currency. Ecology, 73, 1237–1243.

Ashman, T. L., Knight, T. M., Steets, J. A., & Amarasekare, P. (2004). Pollen limitation of plant reproduction: ecological and evolutionary causes and consequences. Ecology, 85, 2408–2421.

Ashmun, J. W., & Pitelka, L. F. (1984). Light-induced variation in the growth and dynamics of transplanted ramets of the understory herb *Aster acuminatus*. Oecologia, 64, 255–262.

Ashmun, J. W., & Pitelka, L. F. (1985). Population biology of *Clintonia borealis*. II. Survival and growth of transplanted ramets in different environments. Journal of Ecology, 73, 185–198.

Barrett, S. C. H. (2015). Influences of clonality on plant sexual reproduction. Proceedings of the National Academy of Sciences of the United States of America, 112, 8859–8866.

Bates, D., Maechler, M., Bolker, B., & Walker, S. (2015). Fitting linear mixed-effects models using lme4. Journal of Statistical Software, 67, 1–48.

Bazzaz, F. A., Chiariello, N. R., Coley, P. D., & Pitelka, L. F. (1987). Allocating resources to reproduction and defense. BioScience, 37, 58–67.

Bennett, E., Roberts, J. A., & Wagstaff, C. (2012). Manipulating resource allocation in plants. Journal of Experimental Botany, 63, 3391–3400.

Bogdanowicz, A., Olejniczak, P., Lembicz, M., & Żukowski, W. (2011). Costs of reproduction in life history of a perennial plant *Carex secalina*. Open Life Sciences, 6, 870–877.

Brunet, J., & Charlesworth, D. (1995). Floral sex allocation in sequentially blooming plants. Evolution, 49, 70–79.

Davis, S. L., & Turner-Jones, L. (2008). Potential for mixed mating in the protandrous perennial *Saponaria officinalis* (Caryophyllaceae). Plant Species Biology, 23, 183–191.

Deevey, E. S. Jr. (1947). Life tables for natural populations of animals. Quarterly Review of Biology, 22, 283–314.

Diggle, P. (1995). Architectural effects and the interpretation of patterns of fruit and seed development. Annual Review of Ecology and Systematics, 26, 531–552.

Diggle, P. K. (1997). Ontogenetic contingency and floral morphology: the effects of architecture and resource limitation. International Journal of Plant Sciences, 158, S99–S107.

Dorken, M. E., van Kleunen, M., & Stift, M. (2025). Costs of reproduction in flowering plants. New Phytologist, 247, 55–70.

Douhovnikoff, V., McBride, J. R., & Dodd, R. S. (2005). *Salix exigua* clonal growth and population dynamics in relation to disturbance regime variation. Ecology, 86, 446–452.

Eckert, C., & Schaefer, A. (1998). Does self-pollination provide reproductive assurance in *Aquilegia canadensis* (Ranunculaceae)? American Journal of Botany, 85, 919–925.

Eriksson, O. (1989). Seedling dynamics and life histories in clonal plants. Oikos, 55, 231–238.

Eriksson, O. (1992). Evolution of seed dispersal and recruitment in clonal plants. Oikos, 63, 439–446.

Eriksson, O. (1999). Seed size variation and its effect on germination and seedling performance in the clonal herb *Convallaria majalis*. Acta Oecologica, 20, 61–66.

Fenner, M. (1985). Seedling establishment. In G. M. Dunnet, & C. H. Gimingham (Eds.), Seed Ecology (pp. 103–116). Springer Netherlands.

Fenner, M. (1987). Seedlings. New Phytologist, 106, 35–47.

Gross, K. L. (1984). Effects of seed size and growth form on seedling establishment of six monocarpic perennial plants. Journal of Ecology, 72, 369–387.

Grime, J. P. (1977). Evidence for the existence of three primary strategies in plants and its relevance to ecological and evolutionary theory. American Naturalist, 111, 1169–1194.

Hamann, E., Wadgymar, S. M., & Anderson, J. T. (2021). Costs of reproduction under experimental climate change across elevations in the perennial forb *Boechera stricta*. Proceedings of the Royal Society B: Biological Sciences, 288, 20203134.

Harper, J. L. (1977). Population Biology of Plants. CAB International.

Hartig, F. (2024). DHARMa: Residual diagnostics for hierarchical (multi-level / mixed) regression models. R package version 0.4.7.

Horvitz, C. C., Ehrlén, J., & Matlaga, D. (2010). Context-dependent pollinator limitation in stochastic environments: can increased seed set overpower the cost of reproduction in an understorey herb? Journal of Ecology, 98, 268–278.

Howe, C. D., & Snaydon, R. W. (1986). Factors affecting the performance of seedlings and ramets of invading grasses in an established ryegrass sward. Journal of Applied Ecology, 23, 139–146.

Ishii, H. S., & Harder, L. D. (2012). Phenological associations of within- and among-plant variation in gender with floral morphology and integration in protandrous *Delphinium glaucum*. Journal of Ecology, 100, 1029–1038.

Jacquemyn, H., Brys, R., & Jongejans, E. (2010). Size-dependent flowering and costs of reproduction affect population dynamics in a tuberous perennial woodland orchid. Journal of Ecology, 98, 1204–1215.

Kliber, A., & Eckert, C. G. (2004). Sequential decline in allocation among flowers within inflorescences: proximate mechanisms and adaptive significance. Ecology, 85, 1675–1687.

Klimeš, L., Klimešová, J., Hendriks, R. J. J., & van Groenendael, J. M. (1997). Clonal plant forms: classification, distribution and phylogeny. In H. Kroon, & J. M. Groenendael (Eds.), The Ecology and Evolution of Clonal Plants. Backhuys Publishers Leiden.

Knight, T. M. (2003). Floral density, pollen limitation, and reproductive success in *Trillium grandiflorum*. Oecologia, 137, 557–563.

Kushwaha, S. P. S., Ramakrishnan, P. S., & Tripathi, R. S. (1983). Population dynamics of *Imperata cylindrica* (L.) Beauv. var. major related to slash and burn agriculture (jhum) in north eastern India. Proceedings of the Indian Academy of Sciences Section B, Biological Sciences, 92, 1–10.

Larson, B. M. H., & Barrett, S. C. H. (1999). The ecology of pollen limitation in buzz-pollinated *Rhexia virginica* (Melastomataceae). Journal of Ecology, 87, 371–381.

Lei, S. A. (2010). Benefits and costs of vegetative and sexual reproduction in perennial plants: a review of literature. Journal of the Arizona-Nevada Academy of Science, 42, 9–14.

Lenth, R. (2025). emmeans: Estimated marginal means, aka least-squares means. R package version 1.11.2-8.

Lokker, C., & Cavers, P. B. (1995). The effects of physical damage on seed production in flowering plants of *Saponaria officinalis*. Canadian Journal of Botany, 73, 235–243.

Medrano, M., Guitián, P., & Guitián, J. (2000). Patterns of fruit and seed set within inflorescences of *Pancratium maritimum* (Amaryllidaceae): nonuniform pollination, resource limitation, or architectural effects? American Journal of Botany, 87, 493–501.

Mitich, L. W. 1990. Bouncingbet—the soap weed. Weed Technology 4, 221–223.

Nilsen, E. T., & Semones, S. (1997). Comparison of variance in quantitative growth and physiological traits between genets and ramets derived from an invasive weed, *Spartium junceum* (Fabaceae). International Journal of Plant Sciences, 158, 827–834.

Obeso, J. R. (2002). The costs of reproduction in plants. New Phytologist, 155, 321–348.

Paz, H., & Martínez-Ramos, M. (2003). Seed mass and seedling performance within eight species of *Psychotria* (Rubiaceae). Ecology, 84, 439–450.

Primack, R. B., & Hall, P. (1990). Costs of reproduction in the pink lady’s slipper orchid: a four-year experimental study. American Naturalist, 136, 638–656.

Qi, S.-S., Dai, Z.-C., Miao, S.-L., Zhai, D.-L., Si, C.-C., Huang, P., Wang, R.-P., & Du, D.-L. (2014). Light limitation and litter of an invasive clonal plant, *Wedelia trilobata*, inhibit its seedling recruitment. Annals of Botany, 114, 425–433.

R Core Team. (2024). R: A language and environment for statistical computing. R Foundation for Statistical Computing.

Rautiainen, P., Koivula, K., & Hyvärinen, M. (2004). The effect of within-genet and between-genet competition on sexual reproduction and vegetative spread in *Potentilla anserina* ssp. *egedii*. Journal of Ecology, 92, 505–511.

Reekie, E. G., & Bazzaz, F. A. (1987). Reproductive effort in plants. I. Carbon allocation to reproduction. American Naturalist, 129, 876–896.

Sandvik, S. M. (2001). Somatic and demographic costs under different temperature regimes in the late-flowering alpine perennial herb *Saxifraga stellaris* (Saxifragaceae). Oikos, 93, 303–311.

Sletvold, N., & Ågren, J. (2015). Climate-dependent costs of reproduction: survival and fecundity costs decline with length of the growing season and summer temperature. Ecology Letters, 18, 357–364.

Sparks, D. (1989). Drought stress induces fruit abortion in pecan. HortScience, 24, 78–80.

Stearns, S. (1989). Trade-offs in life-history evolution. Functional Ecology, 3, 259–268.

Sun, K., Hunt, K., & Hauser, B. (2004). Ovule abortion in *Arabidopsis* triggered by stress. Plant Physiology, 135, 2358–2367.

Therneau, T. M. (2024). coxme: Mixed effects Cox models. R package version 2.2–22.

Totland, Ø. (1997). Limitations on reproduction in alpine *Ranunculus acris*. Canadian Journal of Botany, 75, 137–144.

Tripathi, R. S., & Harper, J. L. (1973). The comparative biology of *Agropyron repens* (L.) Beauv. and A. caninum (L.) Beauv. I. The growth of mixed populations established from tillers and from seeds. Journal of Ecology, 61, 353–368.

Turkington, R. (1979). Neighbour relationships in grass–legume communities. IV. Fine scale biotic differentiation. Canadian Journal of Botany, 57, 2711–2716.

van Noordwijk, A. J., & de Jong, G. (1986). Acquisition and allocation of resources: their influence on variation in life history tactics. American Naturalist, 128, 137–142.

Watkinson, A. R., & Powell, J. C. (1993). Seedling recruitment and the maintenance of clonal diversity in plant populations: a computer simulation of *Ranunculus repens*. Journal of Ecology, 81, 707–715.

Wesselingh, R. A. (2007). Pollen limitation meets resource allocation: towards a comprehensive methodology. New Phytologist, 174, 26–37.

Williams, G. C. (1966). Natural selection, the costs of reproduction, and a refinement of Lack’s principle. American Naturalist, 100, 687–690.

Wu, G.-L., Li, W., Li, X.-P., & Shi, Z.-H. (2011). Grazing as a mediator for maintenance of offspring diversity: sexual and clonal recruitment in alpine grassland communities. Flora, 206, 241–245.

Wulff, R. (1986). Seed size variation in *Desmodium paniculatum*. II. Effects on seedling growth and physiological performance. Journal of Ecology, 74, 99–114.

Zimmerman, M., & Pyke, G. H. (1988). Reproduction in *Polemonium*: assessing the factors limiting seed set. American Naturalist, 131, 723–738.

